# Temporal tuning of repetition suppression across the visual cortex

**DOI:** 10.1101/763078

**Authors:** Matthias Fritsche, Samuel J. D. Lawrence, Floris P. de Lange

## Abstract

The visual system adapts to its recent history. A phenomenon related to this is repetition suppression (RS) - a reduction in neural responses to repeated compared to non-repeated visual input. An intriguing hypothesis is that the timescale over which RS occurs across the visual hierarchy is tuned to the temporal statistics of visual input features, which change rapidly in low-level areas but are more stable in higher-level areas. Here, we tested this hypothesis by studying the influence of the temporal lag between successive visual stimuli on RS throughout the visual system using fMRI. Twelve human volunteers engaged in four fMRI sessions in which we characterized the BOLD response to pairs of repeated and non-repeated natural images with inter-stimulus intervals (ISI) ranging from 50 to 1000 milliseconds to quantify the temporal tuning of RS along the posterior-anterior axis of the visual system. As expected, RS was maximal for short ISIs and decayed with increasing ISI. Furthermore, the overall magnitude of RS gradually increased from posterior to anterior visual areas. Crucially, however, and against our hypothesis, RS decayed at a similar rate in early and late visual areas. This finding challenges the prevailing view that the timescale of RS increases along the posterior-anterior axis of the visual system and suggests that RS is not tuned to temporal input regularities.

## Introduction

How do our brains efficiently process the vast amount of information impinging on our senses? Importantly, sensory input is not random, but exhibits spatial and temporal regularities (Dong and Atick, 1995; Simoncelli and Olshausen, 2001). For instance, our world is relatively stable over short timescales and our brains could exploit this temporal stability in order to optimize neural processing.

One phenomenon that has been linked to optimized processing of visual information over time is repetition suppression (RS) (Muller et al., 1999; Dragoi et al., 2000; Kohn and Movshon, 2004; Grill-Spector et al., 2006; Krekelberg et al., 2006; Gotts et al., 2012). RS refers to the reduction of the neural response to repeated compared to novel sensory input and has been extensively studied and utilized in fMRI research (Tootell et al., 1998; Grill-Spector et al., 1999; Henson et al., 2000; Kourtzi and Kanwisher, 2001; Vuilleumier et al., 2002). Crucially, if RS is a consequence of optimized neural processing due to the exploitation of temporal regularities of input features, it should be adapted to these specific regularities. That is, RS should be limited to timescales for which input features are likely repeated. Importantly however, visual features of different complexity, exhibit different temporal regularities. Simple low-level features, such as contrast edges, represented in primary visual cortex (Hubel and Wiesel, 1959), change rapidly within hundreds of milliseconds, for instance due to eye movements. More complex features, such as faces, represented in downstream ventral areas (Kanwisher et al., 1997), are much more stably present in the input over short timescales. In line with this, it has been proposed that the temporal integration window of neural populations representing increasingly complex features progressively increases along the posterior-anterior axis of the brain (Hasson et al., 2008; Lerner et al., 2011; Honey et al., 2012). The current study was aimed at investigating whether a similar gradient exists for the temporal tuning of RS along the visual hierarchy as measured with fMRI. In particular, we hypothesized that RS should decay over shorter timescales in posterior compared anterior visual areas.

Some indirect evidence for a temporal tuning gradient in RS exists. For instance, previous fMRI studies using brief adaptor stimuli (≤ 1000 ms) have observed RS in primary visual cortex (V1) only for short time lags of 100 to 200 ms between successively repeated input (Kourtzi and Huberle, 2005), but not time lags of 400 ms or longer (Boynton and Finney, 2003; Fang et al., 2005; Kourtzi and Huberle, 2005; Murray et al., 2006; but see Weigelt et al., 2012). Conversely, RS in higher-level object and face selective areas was observed over timescales of several seconds, while these studies generally failed to find RS in early visual areas over similar timescales (Sayres and Grill-Spector, 2006; Weiner et al., 2010). Finally, a recent study reported increasing timescales for the recovery of sub-additive BOLD responses due to repeated visual input along the posterior-anterior axis of the visual system (Zhou et al., 2018).

While these previous findings point towards a gradient in temporal tuning, the evidence is synthesized from studies using different stimuli, timings and tasks, and potentially conflating stimulus-specific with unspecific effects (Huettel and McCarthy, 2001; Zhou et al., 2018). Therefore, in the current study we sought to systematically investigate the short-term temporal decay of RS across the visual hierarchy with one stimulus set consisting of natural images. To preview, we found strong RS to natural images in all visual areas under investigation. Surprisingly, even in V1 RS persisted for temporal lags of 1 second. Furthermore, RS clearly increased along the visual hierarchy, in line with previous findings (Soon et al., 2003; Weiner et al., 2010). Crucially however, against our hypothesis, RS decayed at a similar rate in anterior and posterior visual areas. Therefore, the current results do not provide evidence for the hypothesis that RS reflects a mechanism of optimized neural processing due to the exploitation of temporal input regularities, but are consistent with a more constant mechanism of short-term adaptation throughout the visual system.

## Materials & Methods

### Data availability

All data and code used for stimulus presentation and analysis will be made available on the Donders Institute for Brain, Cognition and Behavior repository (https://data.donders.ru.nl).

### Participants

Fourteen healthy participants (two authors; 9 females, aged 23.4 ± 3.3 years, mean ± SD) took part in the study. All participants reported normal or corrected-to-normal vision and gave written, informed consent prior to the start of the study. The study was approved by the local ethical review board (CMO region Arnhem-Nijmegen, The Netherlands) and was in accordance with the Declaration of Helsinki. Our a priori defined target sample size comprised 12 participants, for which we estimated a power of around 80 to 95% to detect RS effects in visual areas FFA and LOC in response to repeated natural images with an inter-stimulus interval of 500 ms (Kovacs et al., 2013). Participants completed three main task sessions and one session consisting of a retinotopic mapping and functional localizer task. All sessions were conducted on different days. Of the fourteen participants, one participant became unavailable after the first two experiment sessions and one participant was excluded due to excessive head movements. Consequently, data of 12 participants entered data analysis.

### Experimental design

#### Stimuli and experimental paradigm

##### Display

Stimuli were displayed via an LCD projector (EIKI LC-XL100; resolution: 1024 × 768 pixels; refresh rate: 60 Hz) onto a back-projection screen in the bore of the magnet. Participants viewed the screen (distance 96.6 cm; FOV, horizontal 24°, vertical, 18°) through an angled mirror. Stimulus presentation was controlled using the Psychophysics toolbox (Brainard, 1997; Pelli, 1997) for MATLAB (The MathWorks, Natick, MA). For the three main task sessions, eye gaze was monitored and recorded with a high precision Eyelink 1000 eye tracker (SR Research).

##### Main task

An example trial of the main task is depicted in **Fig. 1**. Trials comprised two presentations of natural images (500 ms each), separated by one of five inter-stimulus intervals (50, 100, 200, 500 or 1,000 ms). On a given trial, either the same image could be presented twice (repeat trial) or two different images could be presented (non-repeat trial). The experiment therefore implemented a 2 (repeat/non-repeat) × 5 (ISIs) factorial design. Images were presented in color on a mid-gray background and were windowed with a circular aperture (8° radius) with a linearly ramping boundary (3.2° from 0 to 100%). Each trial was followed by a variable inter-trial interval drawn from a truncated exponential distribution (minimum 2,000 ms; average 3,720 ms). In order to stabilize attention and to avoid potential confounds of attentional factors interacting with ISIs, participants were instructed to perform a demanding one-back digit task at fixation throughout each run (Kay et al., 2013a, 2013b; Zhou et al., 2018). Digits ranging from 0 to 9 (0.25 × 0.25° visual angle) were presented on top of a central, dark gray fixation disk (0.45° diameter). Each digit was on for 500 ms and off for 167 ms before the next digit appeared. Digit presentations were not synchronized to the presentation of the natural images. Participants were asked to press a button every time a digit repeated. On average there were 28 digit repetitions per run (∼ 1 repetition per 30 seconds), and no more than one digit was repeated successively. In order to reduce visual adaptation, the color of digits alternated between black and white for successive presentations. Participants completed four runs (846 seconds each; 1151 volumes, including 10 initial dummy volumes). Each run consisted of 160 trials and contained all ISI levels in pseudo-randomized order. This resulted in 64 trials per repetition x ISI combination per session and thus 192 trials across all three main task sessions. In order to drive visual responses across multiple levels of the visual hierarchy, visual stimuli comprised a set of natural images. Images were taken from the ImageNet database (www.image-net.org). We selected 32 images for each of the categories “Animal”, “Body”, “Face’”, “Fruit”, “House”, “Indoor”, “Landscape”, “Object”, “Outdoor” and “Vehicle”, resulting in a total of 320 images. For each participant and session, we created 160 randomly assigned pairs of images A and B from the full set of 320 images, under the constraint that A and B were of different categories. Image pairs were then randomly assigned to one of the five ISI conditions. Each image pair was presented once per run in one of the four possible orders (A-A, B-B, A-B or B-A), and every order was presented once per session. Consequently, the overall visual stimulation across repeats and non-repeat trials within each ISI condition was equal.

**Figure 1.**
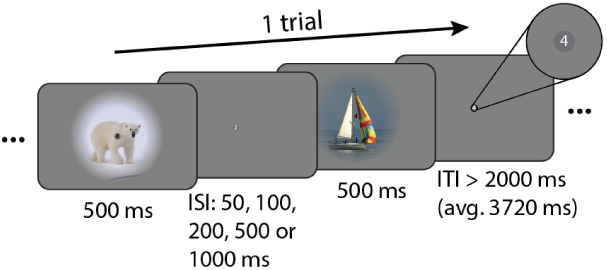
Example trial of the main task. Each trial comprised two stimulus presentations of 500 ms each. Either the same image was presented twice (repeat trial) or two different images were presented (non-repeat trial). Image presentations were separated by one of five inter-stimulus intervals (50, 100, 200, 500 or 1000 ms). Participants were instructed to fixate on a small gray disk in the center of the screen (see inset) and performed a 1-back task on digits (0 to 9), which were shown within the fixation disk at a presentation rate of 1.5 Hz. Trials were separated by a variable inter-trial-interval, during which only the fixation disk and digits were shown.

##### Functional localizer

For mapping functional ROIs, participants viewed centrally presented greyscale images (16 × 16° visual angle) from the fLoc functional localizer package (Stigliani et al., 2015), while fixating on a central fixation dot (0.2° visual angle diameter). Stimuli were presented in a blocked design. Each block lasted for 16 seconds and comprised 32 individual images of the same category (2 Hz presentation rate). Each run (423 seconds; 575 volumes, including 10 initial dummy volumes) consisted of 26 blocks, of which the first and last blocks were always baseline blocks in which no images were presented. The 24 interjacent blocks consisted of object, scene, face, body part, phase-scrambled and baseline blocks with the same frequency (four per each run). The order of blocks was pseudorandomized such that adjacent blocks were always of a different category. Participants performed a 1-back task on the images. Each participant completed three runs in total.

##### Retinotopy

For retinotopic mapping, participants viewed high contrast, contrast-reversing (6 Hz) 90° rotating wedge and expanding ring stimuli that mapped responses to the central region of the visual field (radius 9 degrees of visual angle). Stimulus runs lasted 297 s (198 volumes), including eight full stimulus cycles and six starting dummy volumes that were discarded prior to analyses. During stimulus runs, participants were instructed to fixate on a small central fixation cross (0.24° visual angle across) and press a button every time it changed color from red to green or green to red (30 switches per run). Participants completed six runs of wedge stimulus presentation and one run of ring stimulus presentation.

##### fMRI data acquisition

Functional and anatomical images were collected on a 3T Prisma Fit MRI system (Siemens), using a 32-channel headcoil. Functional images in the main task and functional localizer were acquired using a whole-brain T2*-weighted multiband-6 sequence (TR/TE = 735/39 ms, 48 slices, voxel-size 2.4 mm isotropic, 52° flip angle, A/P phase encoding direction). Functional images in the retinotopy task were acquired using a whole-brain T2*-weighted multiband-4 sequence (TR/TE = 1500/39.6 ms, 68 slices, voxel-size 2 mm isotropic, 75° flip angle, A/P phase encoding direction). One high resolution anatomical image was acquired with a T1-weighted MP-RAGE sequence (TR/TE = 2300/3.03 ms, voxel-size 1 mm isotropic, 8° flip angle, GRAPPA acceleration factor 2).

### Data analysis

#### fMRI data preprocessing

fMRI data of the main and functional localizer tasks were preprocessed using FSL 6.00 (FMRIB Software Library; Oxford, UK; www.fmrib.ox.ac.uk/fsl; Smith et al., 2004). The preprocessing pipeline encompassed brain extraction (BET), motion correction (MCFLIRT), temporal high-pass filtering (100 s cutoff), spatial smoothing with a Gaussian kernel (FWHM of 5 mm) and low-pass filtering using a Savitzky-Golay filter with window length m = 7 TRs and polynomial order n = 3 (Savitzky and Golay, 1964). The first 10 volumes of each run were discarded to allow for signal stabilization.

#### Main task analysis

To estimate BOLD activations to repeat and non-repeat trials at different ISIs, we employed finite impulse response (FIR) models (Ollinger et al., 2001). Briefly, we defined FIR basis functions of unit impulse from 0 to 16 seconds after trial onset, in steps of 1 seconds for each of the 10 trial types (5 ISIs x repeat/non-repeat). This resulted in a total of 170 regressors (17 timepoints x 10 trial types), which were used to fit voxelwise GLMs to each participant’s run data using FSL FEAT. Subsequently, the estimated BOLD responses for each trial type were averaged across runs and across all voxels within a given ROI (see *Functional ROI definition & Retinotopic ROI definition*) and fit with a double gamma function of the form

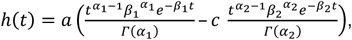

where *t* designates time from trial onset, *a* controls the amplitude, α and β control the shape and scale, respectively, and *c* determines the ratio of the response to undershoot (Lindquist et al., 2009). The peaks of the positive response and late undershoot were constrained to occur between 3 and 8 seconds, and 8 and 18 seconds, respectively (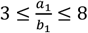 and 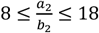). Given the current experimental design, this approach of estimating the shape of the hemodynamic response function (HRF) from the data appears more applicable than a standard GLM analysis assuming a single canonical HRF for all trial types and brain areas. First, due to the different ISIs across trials, the onset and peak latencies of the BOLD response are expected to differ slightly across ISI conditions (see **Supplemental Fig. S1**). Second, it is likely that HRFs will differ in shape across regions. Both aspects could lead to biased activation estimates across ISIs and ROIs in a standard GLM. From the estimated models, we derived the peak amplitudes *A* for each trial type (**Fig. 2**). Subsequently, we quantified RS within each ROI at every ISI as

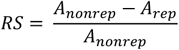

where *A*_*rep*_ and *A*_*nonrep*_ reflect the peak BOLD amplitudes to repeat and non-repeat trials, respectively. In other words, the *RS* score denotes the percentage reduction of BOLD amplitude for repeat relative to non-repeat trials. Of note, the *RS* score captures the BOLD amplitude reduction to the compound BOLD response of the first and second image of each trial, and therefore does not directly reflect the response reduction to the second repeated or non-repeated image alone. Since the BOLD response to the first image will be similar for repeat and non-repeat trials, the actual response reduction to the second image will be larger than indicated by the *RS* score, which factors in responses to both images. In fact, for the ISIs employed in this experiment, the actual response reduction to the second image will be approximately double the size of the estimated *RS* score. That is, a *RS* score of 25% corresponds to a response reduction of ∼50% to the second repeated versus non-repeated image. Importantly, simulations reveal that the relationship between *RS* scores and the corresponding response reduction to the second image are preserved across the different ISIs employed in this experiment, allowing an unbiased comparison RS across the different ISIs (see **Supplemental Fig. S1**). To estimate the decay of RS with increasing ISI, we fit linear models to each participant’s *RS* scores as a function of ISI, such that the slope of the model described the change of *RS* scores over time. Intercepts and slopes of these linear models were statistically evaluated with separate repeated measures ANOVAs. We applied Greenhouse-Geisser correction, when a violation of sphericity was indicated. We hypothesized to find reduced BOLD amplitudes for repeat versus non-repeat trials, leading to positive model intercepts. Furthermore, we expected RS to decay with increasing temporal lag between stimulus repetitions, which should be reflected in overall negative slopes. To assess the hypothesis of a faster decay of RS for posterior compared to anterior visual areas we sought to compare the decay of RS across areas independently of the initial magnitude of RS in each area. To this end, we first normalized each participant’s slope estimate of a given ROI by its corresponding group mean intercept estimate. Consequently, the normalized slope parameter expressed the decay of RS relative to the initial magnitude of RS within a visual area and thus accounts for differences in slope due to differences in the initial magnitude of RS across visual areas. The normalized slope estimates were statistically evaluated with a repeated measures ANOVA. We expected to find a faster decay of RS for posterior compared to anterior visual areas, which would be captured by steeper, more negative slopes for more posterior compared to anterior ROIs. To test the hypothesis of gradually steeper slopes for posterior compared to anterior ROIs more directly, we further computed Spearman rank correlations between the normalized RS slope estimates and the position of the ROIs in the visual hierarchy. While areas V1 to V3 were assigned to successively increasing levels in the visual hierarchy (Felleman and Van Essen, 1991), we assigned areas V4, LO1, LO2 and LOC to the same hierarchical level, as there is no clearly established distinction in hierarchical level between ventral V4 and dorsal LO1/2, and as the retinotopically defined areas LO1/2 and functionally defined area LOC are partly overlapping (Larsson and Heeger, 2006). Finally, areas FFA and PPA were assigned the highest level among the ROIs of the current experiment. The rank-order correlation assessed a monotonic relationship between an area’s position in the visual hierarchy and its rate of decay of the RS effect. Rank-order correlations were computed for each individual participant and the resulting correlation coefficients were tested against zero on the group-level using a two-sided Wilcoxon signed-rank test, with a significance threshold of α = 0.05. We expected to find a positive correlation between the ROIs’ position in the visual hierarchy and their normalized RS slope estimates, indicating a gradually slower decay of RS in higher-level visual areas.

**Figure 2.**
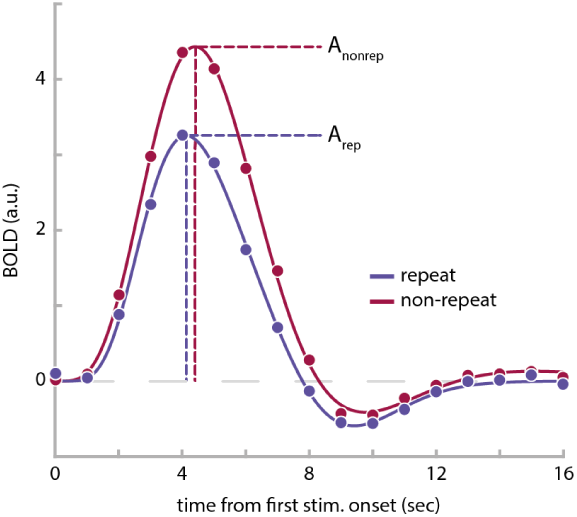
Estimated BOLD responses in area PPA of one example observer for trials with an inter-stimulus interval of 50 ms. The BOLD response for repeat trials (blue data points) is reduced in comparison to non-repeat trials (red data points). BOLD amplitudes, A_rep_ and A_nonrep_, were estimated by fitting double-gamma functions (solid lines) to the BOLD time courses and extracting the peak height of the double gamma function, respectively. See **Supplemental Fig. S2** for group averaged BOLD responses and their respective double-gamma model fits for each ROI and ISI combination.

#### Functional ROI definition

In order to map face, place and object selective brain areas, voxelwise GLMs were fit to each participant’s run data using FSL FEAT. Separate regressors for object, scene, face, body part, phase-scrambled and baseline blocks were added to the model. Blocks were modeled with a 16 seconds duration boxcar and convolved with a gamma hemodynamic response function. Moreover, nuisance regressors in the form of first-order temporal derivatives of all event types and 24 motion regressors (six motion parameters, the derivatives of these motion parameters, the squares of the motion parameters, and the squares of the derivatives; comprising FSL’s standard extended set of motion parameters) were added to the model. Data were combined across runs using FSL’s fixed effect analysis. For identifying object-selective area LOC, we contrasted average activations to objects versus scrambled blocks. For identifying face-selective area FFA and place-selective area PPA we contrasted activations of faces versus scene blocks, and scene versus face blocks, respectively. Differential activations were visualized on an inflated cortical surface and initially thresholded at *p* < 1e-4 (uncorrected). When necessary, the thresholds were raised on a per participant and per ROI basis until clusters of activations where limited to a reasonable extent, comparable in size and location to previous studies (Epstein and Kanwisher, 1998; Grill-Spector et al., 2001; Kanwisher and Yovel, 2006; Sayres and Grill-Spector, 2006).

#### Retinotopic ROI definition

Anatomical data were automatically segmented into white matter, gray matter and CSF using FreeSurfer (http://surfer.nmr.mgh.harvard.edu/) and manually corrected. Functional data were analyzed using the phase encoding approach in MrVista (http://white.stanford.edu/software/). Polar angle and eccentricity data were visualized on an inflated cortical surface and the boundaries of V1, V2 and V3, V4, LO1 and LO2 were drawn manually using established criteria (Engel et al., 1994; Sereno et al., 1995; Larsson and Heeger, 2006; Wandell et al., 2007; Amano et al., 2009; Winawer and Witthoft, 2017).

## Results

Repeating visual stimuli led to a clear reduction of BOLD amplitude across all visual areas (**Fig. 3A**). Accordingly, linear model fits to *RS* scores of each individual *ROI* exhibited significantly positive intercepts, indicating robust RS (*F*(1,11) = 92.5, p = 1e-6). Furthermore, RS gradually increased from posterior to anterior visual areas (**Fig. 3A**), supported by a significant effect of ROI on the intercept term of the linear models (**Fig. 3B**, *F*(2.5,27.3) = 6.2, p = 0.004). This was further corroborated by a significant positive correlation between the ROI position in the visual hierarchy and the intercept of RS (mean rho = 0.63 ± 0.10 SEM; Wilcoxon W = 76, p = 0.004). In fact, on the individual participant level, 8 out of 12 participants showed a significant positive correlation (p < 0.05). Moreover, in line with prior research (Henson et al., 2000, 2004), the strength of RS decreased with increasing temporal lag between repeated stimuli, as revealed by consistently negative model slopes (**Fig. 3C**, *F*(1,11) = 11.6, p = 0.006). Crucially however, our initial hypothesis that posterior visual areas would show a steeper decay of RS compared to anterior areas was not borne out in the data. First, a repeated-measures ANOVA revealed no significant effect of ROI on normalized slopes (**Fig. 4**, *F*(2.7,29.3) = 0.33, p = 0.78). Second, a more sensitive rank-order correlation analysis revealed no significant correlation between the ROI position in the visual hierarchy and the normalized slope of the RS decay (mean rho = - 0.21 ± 0.18 SEM; Wilcoxon W = 15, p = 0.23). Instead, RS decay was relatively similar across regions, such that RS was reduced to approximately half of its initial magnitude in all areas after 1000 ms (**Fig. 4**). Interestingly, this meant that RS was observed for a prolonged period of time even in the earliest regions of visual cortex. Against our expectations, V1 showed robust RS throughout all temporal lags, even for the longest lag of 1000 ms (*t*(11) = 2.9, p = 0.016). This stands in apparent contrast to several previous findings on short-term adaptation, which do not find stimulus-specific RS in early visual cortex for temporal lags exceeding a couple hundred of milliseconds (Boynton and Finney, 2003; Kourtzi and Huberle, 2005; Murray et al., 2006; Weiner et al., 2010).

**Figure 3.**
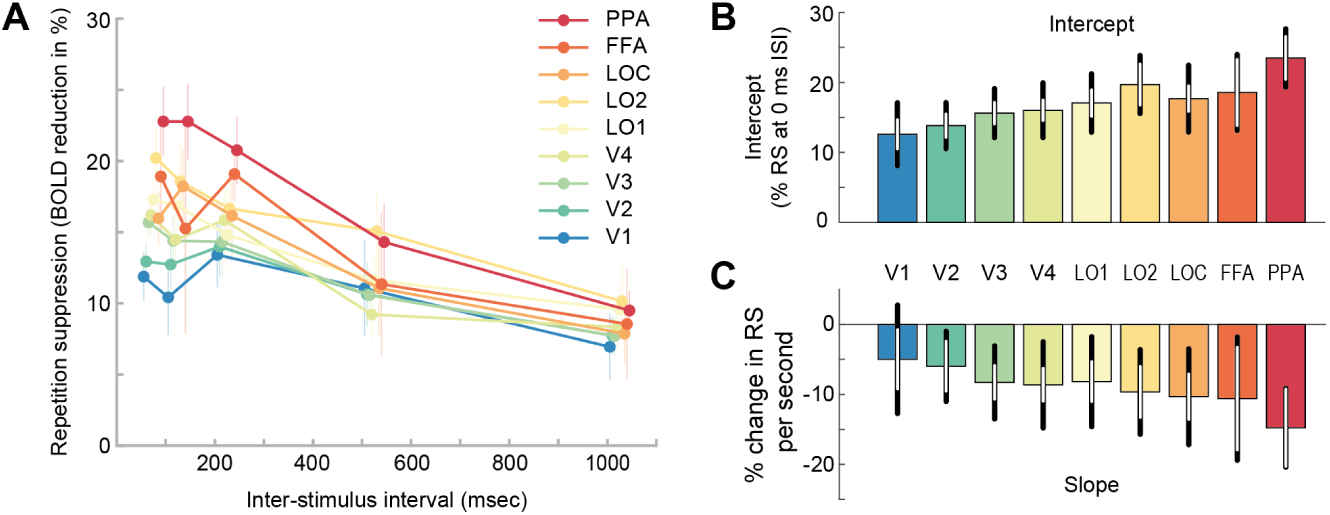
Repetition suppression across visual areas and inter-stimulus intervals. **A**, Repetition suppression (*RS*) score of each ROI (color-coded) as a function of inter-stimulus interval. The *RS* score indicates the percentage reduction of BOLD amplitude for repeat relative to non-repeat trials. Positive *RS* scores mark RS. While all ROIs show repetition suppression, the strength of RS increases from posterior to anterior regions. Furthermore, RS decreases for increasing temporal lags between image presentations. Error bars denote SEMs. **B**, Mean intercept estimates of linear models fitted to single subject data in **A**. Intercept estimates increase from posterior to anterior regions. Black error bars denote 95% confidence intervals. White error bars denote 95% within-subject confidence intervals. **C**, Mean slope estimates of linear models, indicating the decay of *RS* scores over time. See **Supplemental Fig. S3** for single subject estimates of intercepts and slopes, **Supplemental Fig. S4** for model fits and **Supplemental Fig. S5** for residual plots.

**Figure 4.**
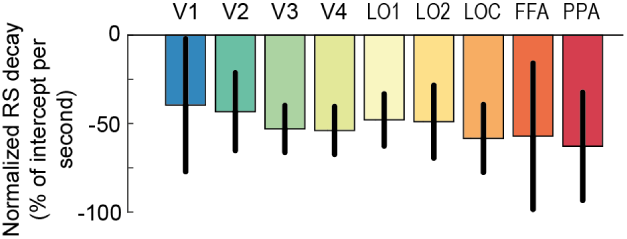
Decay of repetition suppression normalized to its initial magnitude. The slope parameter estimates of the linear model fits to *RS* scores (**Fig. 3C**) are normalized by the group mean intercept estimates for each region (**Fig. 3B**), respectively. The normalized slope parameter estimate accounts for differences in slope due to differences in the initial magnitude of RS across visual areas. RS decays similarly across posterior and anterior visual areas and is reduced to approximately half of its initial magnitude after 1 second. Error bars denote 95% within-subject confidence intervals.

## Discussion

The aim of the current study was to investigate the temporal tuning of short-term RS to natural images across the visual system. Overall, we found strong reductions in BOLD responses for repeated compared to non-repeated images across all visual areas under investigation. Furthermore, we found a gradual increase in the strength of RS from posterior to anterior regions and a gradual decay of RS with increasing temporal lag between stimuli. Crucially, however, against our initial hypothesis, there was no evidence for a faster decay of RS for low-compared to high-level visual areas. Instead, RS in low- and high-level visual areas appeared to decay at a similar rate. Therefore, the current results do not provide evidence for the hypothesis that short-term RS in the visual system is tuned to the temporal statistics of visual features of different complexity, and challenge a prevailing view that RS generally persists over longer timescales in high-level cortical areas (Barron et al., 2016).

The finding that RS does not decay faster in posterior compared to anterior visual areas stands in apparent contrast to several previous findings, which collectively suggest such a differential decay. For instance, previous studies demonstrated RS in higher-level object selective cortex over time-periods of several seconds, in the absence of similar effects in early visual areas (Sayres and Grill-Spector, 2006; Weiner et al., 2010). Notably, RS in higher-level occipitotemporal regions has been observed over time lags of minutes (Henson et al., 2000, 2004) and days (van Turennout et al., 2000, 2003). In contrast, RS in primary visual cortex is typically only observed for very short temporal lags of around 200 ms (Kourtzi and Huberle, 2005; Kourtzi et al., 2010), but absent for longer lags (Boynton and Finney, 2003; Fang et al., 2005; Kourtzi and Huberle, 2005; Murray et al., 2006), although RS for lags of 1 second has been found in long-term adaptation designs with initial adaptors of 100 seconds duration and 4 seconds long top-up adaptors (Larsson et al., 2006; Montaser-Kouhsari et al., 2007). Our current findings differ from these previous reports in two aspects. First, in the current experiment RS in anterior object-selective cortex decreases rapidly over the tested time window, at a rate of about 50% of its initial magnitude per second. Extrapolating this to longer time windows, one would expect that RS should be completely absent for lags longer than ∼2 seconds. Second, we find sustained RS to brief adaptors in early visual areas, such as V1, for temporal lags far longer than 200 ms. In fact, there is robust RS in V1, even at a temporal lag of 1 second.

What could account for the substantially faster decay of RS in anterior regions, compared to previous studies? In contrast to most previous studies on RS, which employed tasks that required participants to actively attend and process each stimulus, the current experiment involved a demanding task at fixation, which diverted attention away from the repeated or non-repeated visual stimulation. Attention has been shown to modulate RS effects (Murray and Wojciulik, 2004; Henson and Mouchlianitis, 2007; Larsson and Smith, 2012) and it is conceivable that it may change the temporal dynamics of RS across cortical areas. That is, attending and actively processing visual stimuli might lead to long-term synaptic changes in higher-level cortical areas, which could support reductions of neural responses over longer timescales of minutes to days. Related to this, long-term RS involving several intervening stimuli between repetitions appears to be influenced by the precise task performed on the stimuli (Henson et al., 2002) and stimulus familiarity (Henson et al., 2000). Generally, it seems likely that long-term RS effects in category-selective areas reflect a combination of memory-dependent and task-dependent processes (see Henson, 2016), which are very different from the process of short-term sensory adaptation studied in the current experiment. In turn, the current results indicate that, while short-term RS can be observed in category-selective cortex in the absence of attention towards the stimuli, it is characterized by a much more rapid temporal decay than suggested by previous findings.

Intriguingly, in contrast to previous studies, we also observed robust RS to brief adaptor stimuli in V1 for temporal lags of up to 1000 ms. One clear difference between the current and previous studies is the stimulus set used to probe RS. While previous studies examining short-term adaptation in early visual cortex typically employed simple grating stimuli (Boynton and Finney, 2003; Fang et al., 2005; Murray et al., 2006) or 2-D shape stimuli consisting of locally oriented Gabors (Kourtzi and Huberle, 2005), the current study used natural images to drive visual responses across multiple levels of the visual hierarchy. Importantly, previous electrophysiological studies have shown differences in the oscillatory neural activity elicited by artificial grating stimuli versus naturalistic visual input in early visual areas, likely due to differences in both the temporal and spatial input statistics (Kayser et al., 2003; Hermes et al., 2015). In particular, in contrast to many natural images, grating stimuli elicit increased narrowband gamma power (Hermes et al., 2015), which has been linked to gain control or normalization (Ray et al., 2013; Hermes et al., 2019). If these inhibitory normalization processes, which are strongly engaged by grating stimuli, are themselves subject to adaptation, this may explain the generally weaker and less sustained RS effects for grating stimuli in comparison to natural images. While this explanation would mainly rely on adaptation of inhibitory lateral connections within V1, there is an alternative explanation involving recurrent connections across cortical areas. Importantly, since BOLD responses reflect an integral of neural activity over several seconds, it is possible that the RS effect observed in V1 at a lag of 1000 ms is due to feedback signals from higher-level areas, which maintain RS over longer timescales. If natural images elicit more recurrent activity across the visual hierarchy than grating stimuli, such an indirect RS effect due to feedback signals from high-level visual areas could explain the different lag dependence in V1. Future studies may directly compare short-term RS effects for gratings and natural stimuli and assess potential effects of recurrent processing with electrophysiological methods and laminar fMRI.

In apparent contradiction to the current results, a recent study has found that early visual areas show a faster recovery from sub-additive BOLD responses due to repeated visual input in comparison to downstream areas (Zhou et al., 2018). While BOLD responses to two stimuli presented in rapid succession were smaller than predicted by the sum of BOLD responses to an individual stimulus, the BOLD response in early visual cortex quickly approached the linear prediction for increasing temporal lags between stimulus presentations. Conversely, BOLD responses in more anterior visual areas remained below their linear prediction over progressively longer timescales. This adaptation effect and its temporal tuning across visual areas was adequately captured by a temporal compressive summation model. However, it should be noted that Zhou et al. studied adaptation using repeated presentations of the same visual stimulus, and therefore cannot distinguish between stimulus-specific and unspecific adaptation. Crucially, stimulus-unspecific adaptation effects have been widely observed in Fmri studies investigating the effects of stimulus repetition (Boynton and Finney, 2003; Soon et al., 2003; Kourtzi and Huberle, 2005; Murray et al., 2006) and are likely also present in the data by Zhou et al. (2018). Crucially, such unspecific adaptation effects may be driven by vascular mechanisms, unrelated to neural adaptation, and therefore it is important to separate them from stimulus-specific adaptation. By contrasting BOLD responses of repeat and non-repeat trials, for which the overall visual input was equated, we account for stimulus-unspecific adaptation effects in our analysis.

Furthermore, we have found that RS gradually increased in magnitude along the posterior-anterior axis of the visual system. There exist several explanations for such a gradient. First, it is possible that anterior regions exhibit intrinsically stronger RS than posterior regions. Second, RS effects in downstream cortical regions could, to some degree, be inherited from upstream regions (Kohn and Movshon, 2003; Larsson and Harrison, 2015). Such an inheritance effect would naturally produce an incremental increase of RS along the visual hierarchy. Lastly, increased RS in area MT compared to V1 has been attributed to an interaction of receptive field and stimulus size, leading to different amounts of surround suppression across areas (Wissig and Kohn, 2012; Patterson et al., 2014). However, these findings were based on drifting grating stimuli and only occurred for prolonged adaptation of 40 seconds, but not short adaptation used in the current study.

Finally, we would like to point out two important limitations of the current study. First, our current results may depend on the stimulus set and experiment parameters such as adaptor and test stimulus timing employed in this study. As mentioned above, using grating stimuli instead of natural images may change the strength and temporal dynamics of RS across visual areas. For instance, different stimulus families, such as gratings versus natural images, may drive the neural circuitry, which ultimately gives rise to adaptation effects, in different ways. Furthermore, the duration of the adaptor stimulus will likely influence the magnitude and decay of RS and these effects could potentially differ across visual areas. For instance, while RS in V1 to brief adaptor gratings (≤ 1 second) appears to be absent for temporal lags longer than 400 ms (Boynton and Finney, 2003; Fang et al., 2005; Kourtzi and Huberle, 2005; Murray et al., 2006), RS effects for a lag of 1 second have been found in long-term adaptation designs, where the adaptor grating is presented for 100 seconds (Larsson et al., 2006; Montaser-Kouhsari et al., 2007). In the current study we were interested in whether the timescale over which RS occurs across the visual hierarchy is tuned to the temporal statistics of visual input features. Therefore, we chose to focus on short-term adaptation to natural images, containing a wide range of input features of differing complexity and approximating natural viewing conditions. Extending the current results to other stimulus sets, stimulus timings and tasks is an interesting topic for future research. A second limitation derives from the observation that adaptation effects in downstream cortical regions can be inherited from upstream regions (Kohn and Movshon, 2003; Larsson and Harrison, 2015). The potential presence of such inheritance effects may complicate the assessment of intrinsic temporal dynamics of RS within a particular visual area, independently of the dynamics in upstream areas. While feedforward inheritance effects can to some extent be circumvented by changing the spatial location of adaptor and test stimuli, thus exploiting the increase in receptive field size from low- to high-level visual areas (Larsson and Harrison, 2015), this would preclude a direct comparison of adaptation in low and high-level visual areas. Furthermore, inheritance effects might not be limited to feedforward processing, as adaptation effects in higher-level areas could be passed down to lower-level regions via recurrent connections. When interpreting the results of the current study, it should be noted, however, that if the temporal decay of RS in higher-level regions would be mainly driven by the decay of RS in lower-level regions, and higher-level regions would add little to no RS themselves, such a pattern RS across the visual cortex would be incompatible with a view of optimized visual processing throughout the visual hierarchy, in which each area would exploit the temporal regularities of the features which it is tuned to.

To conclude, the current study revealed strong RS to natural images across the visual hierarchy, even in primary visual cortex, up to delays of 1,000 ms. Crucially however, RS does not appear to decay more slowly in anterior compared to posterior visual areas. Thus, the current results do not provide evidence for the hypothesis that short-term RS is tuned to the temporal statistics of visual features with different complexities. This suggests that RS is not a consequence of optimized neural processing due to the exploitation of temporal regularities of input features, but appears to reflect an adaptation process that has a fixed time constant across the visual system. This may limit the specificity with which the mechanisms underlying RS could optimize the processing of input features at any level of the visual hierarchy.

## Acknowledgements

All authors designed the study. M.F. and S.J.D.L. programmed the stimuli. M.F. and S.J.D.L. collected the data. M.F. analyzed the data. All authors wrote the manuscript. This work was supported by a grant from the European Union Horizon 2020 Program (ERC Starting Grant 678286, “Contextvision”).

## Supplemental data

**Supplemental Fig. S1.**
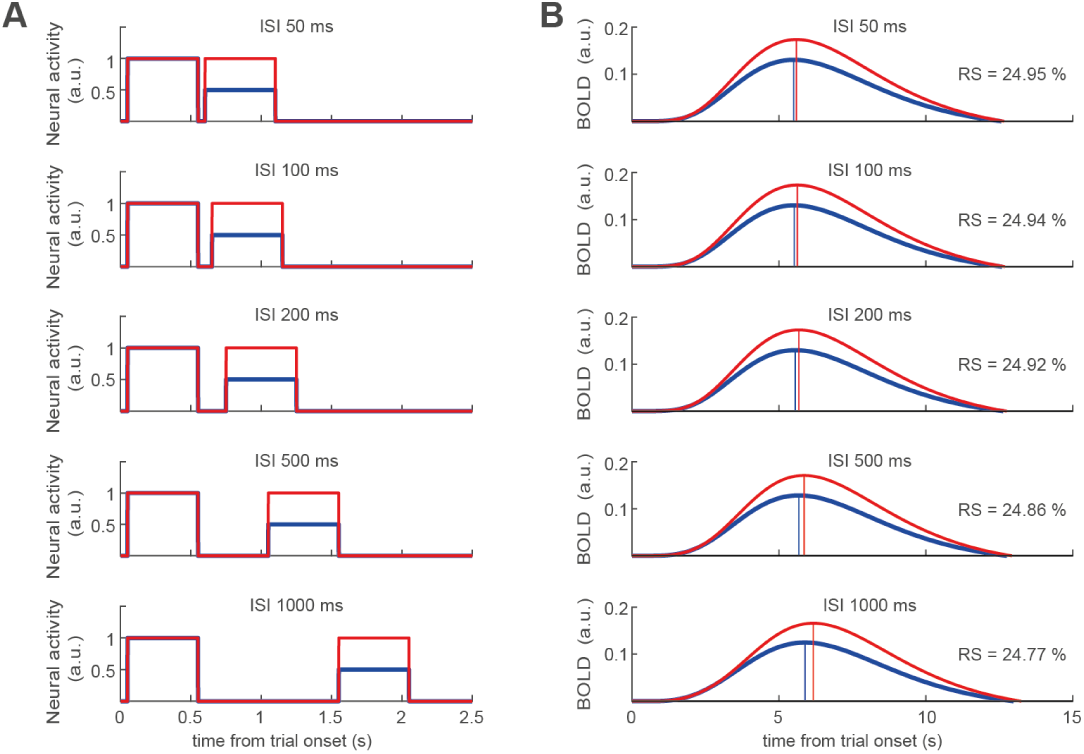
Relationship between neural activations and BOLD response across different inter-stimulus intervals. Due to the slow dynamics of the BOLD signal, the BOLD responses measured in the current experiment reflect compound responses to individual neural activations elicited by two stimuli presented in rapid succession within one trial. For instance, when modeling neural activations to individual images of 500 ms duration as boxcar functions of the same duration (**panel A**), and convolving two such activations occurring in rapid succession with a standard double-gamma hemodynamic response function, one obtains unimodal BOLD responses (**panel B**). Since the BOLD response is driven by neural activations to both the first and second stimulus, a 50% reduction of neural activation to a second repeated stimulus (blue traces in **A**) relative to a non-repeated stimulus (red traces in **A**) only translates into a ∼25% relative reduction of the corresponding BOLD responses (blue and red traces in **B**). Therefore, the empirically measured *RS* scores likely reflect reductions in neural activations that are approximately twice the size of their corresponding *RS* scores. Furthermore, one might be worried that the relationship between *RS* scores and underlying reductions in neural activity is influenced by the varying inter-stimulus interval (**panel A**, top to bottom). While they overall BOLD amplitude indeed decreases with increasing ISIs, even though the overall amount of neural activations remains constant, the relative reduction of BOLD amplitudes for repeat versus non-repeat trials remains approximately constant (see panel B, inset values of RS scores). This allows to meaningfully relate changes in *RS* scores across different ISIs to changes in underlying neural activations.

**Supplemental Fig. S2.**
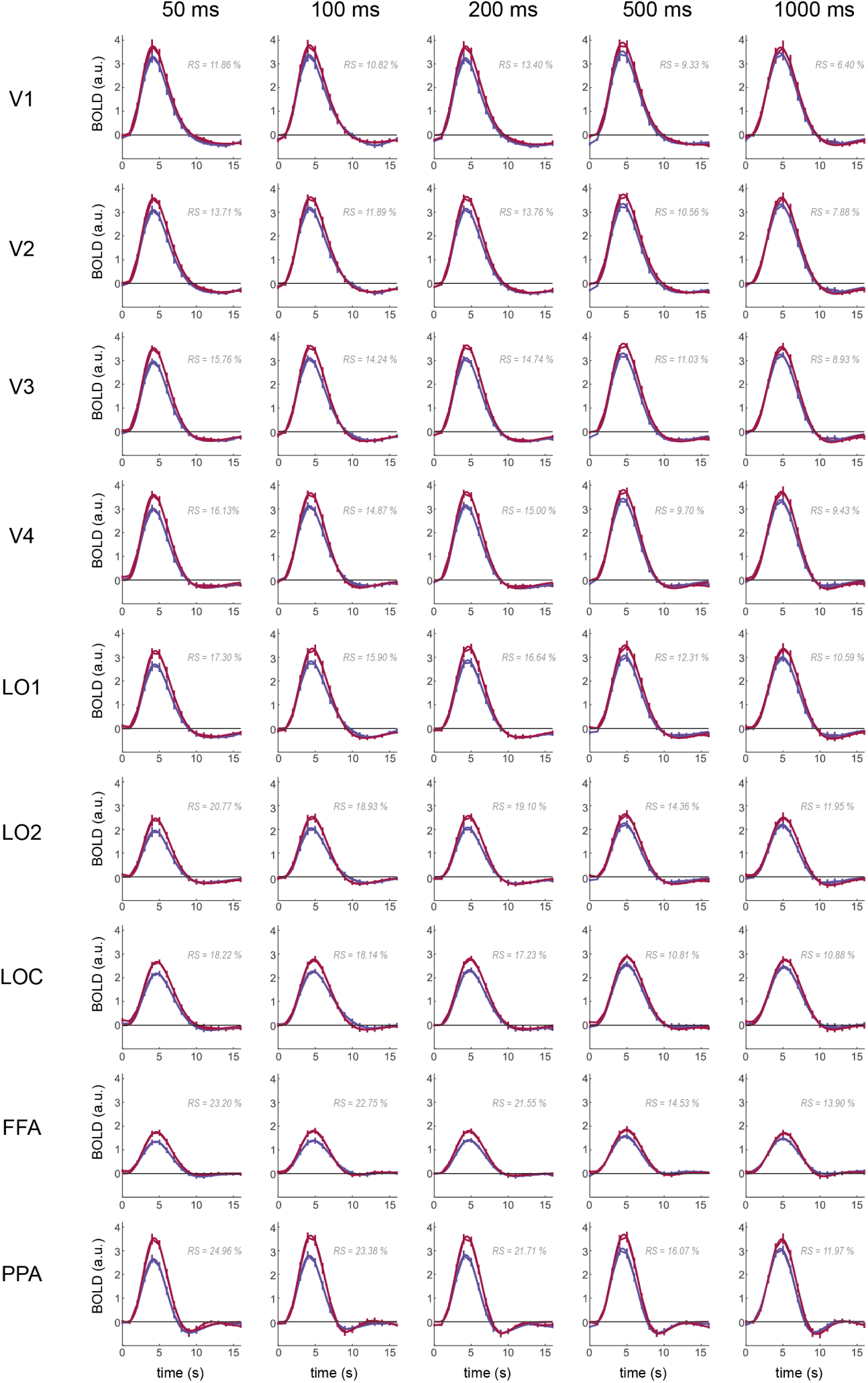
Group average BOLD responses for repeat (blue) and non-repeat (red) trials for each ROI (rows) and ISI (columns). Thick lines show double-gamma model fits. Gray numbers indicate the *RS* score, i.e. the percentage reduction of BOLD amplitude for repeat relative to non-repeat trials, as computed from the model fits to the group average BOLD responses. Note that these *RS* scores numerically slightly deviate from the mean *RS* scores in Fig. 3A, since those scores were computed on single-subject BOLD estimates and subsequently averaged. Error bars denote SEMs.

**Supplemental Fig. S3.**
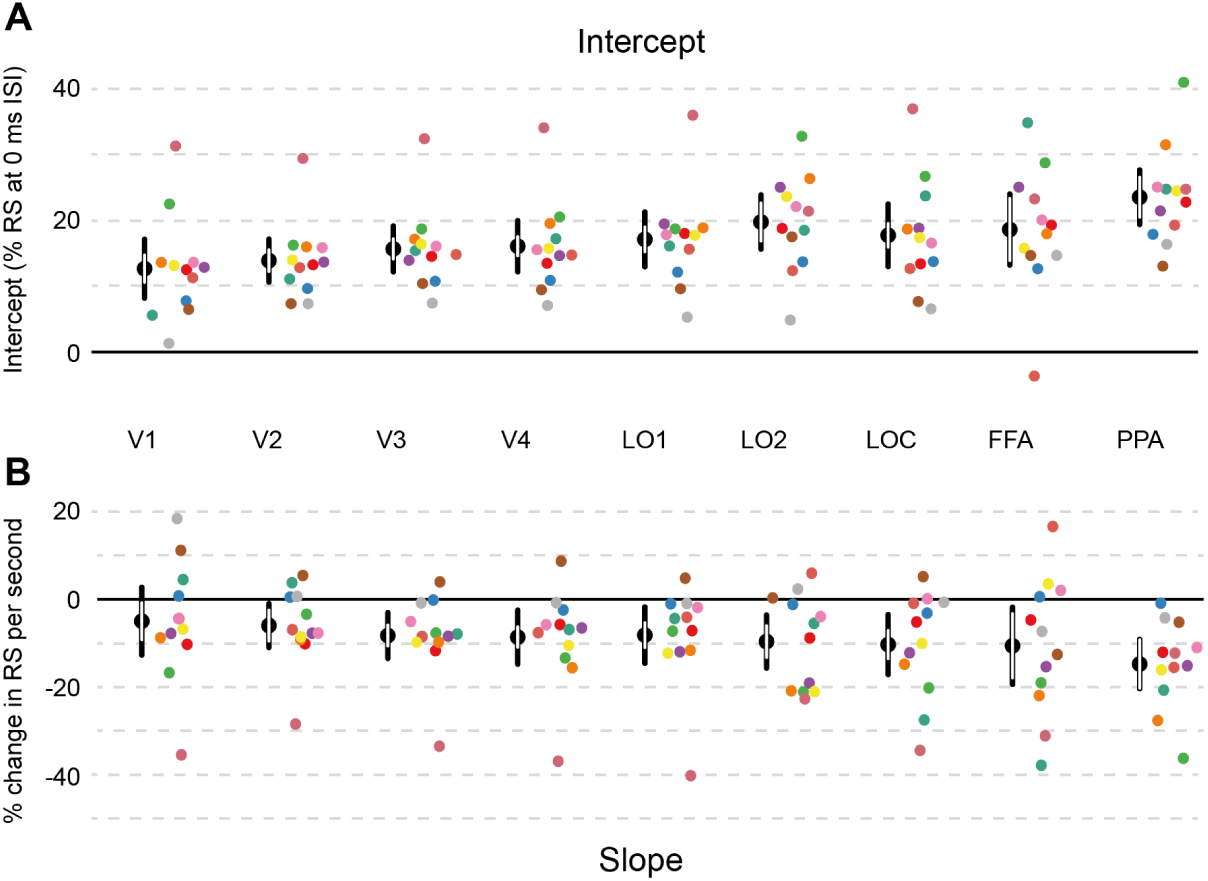
Intercept (**A**) and slope (**B**) estimates of linear models fitted to single subject data in **Fig. 3A** (see also **Supplemental Fig. S4**). Colored data points show single subject estimates, while black data points show the mean parameter estimate. Black error bars denote 95% confidence intervals. White error bars denote 95% within-subject confidence intervals.

**Supplemental Fig. S4.**
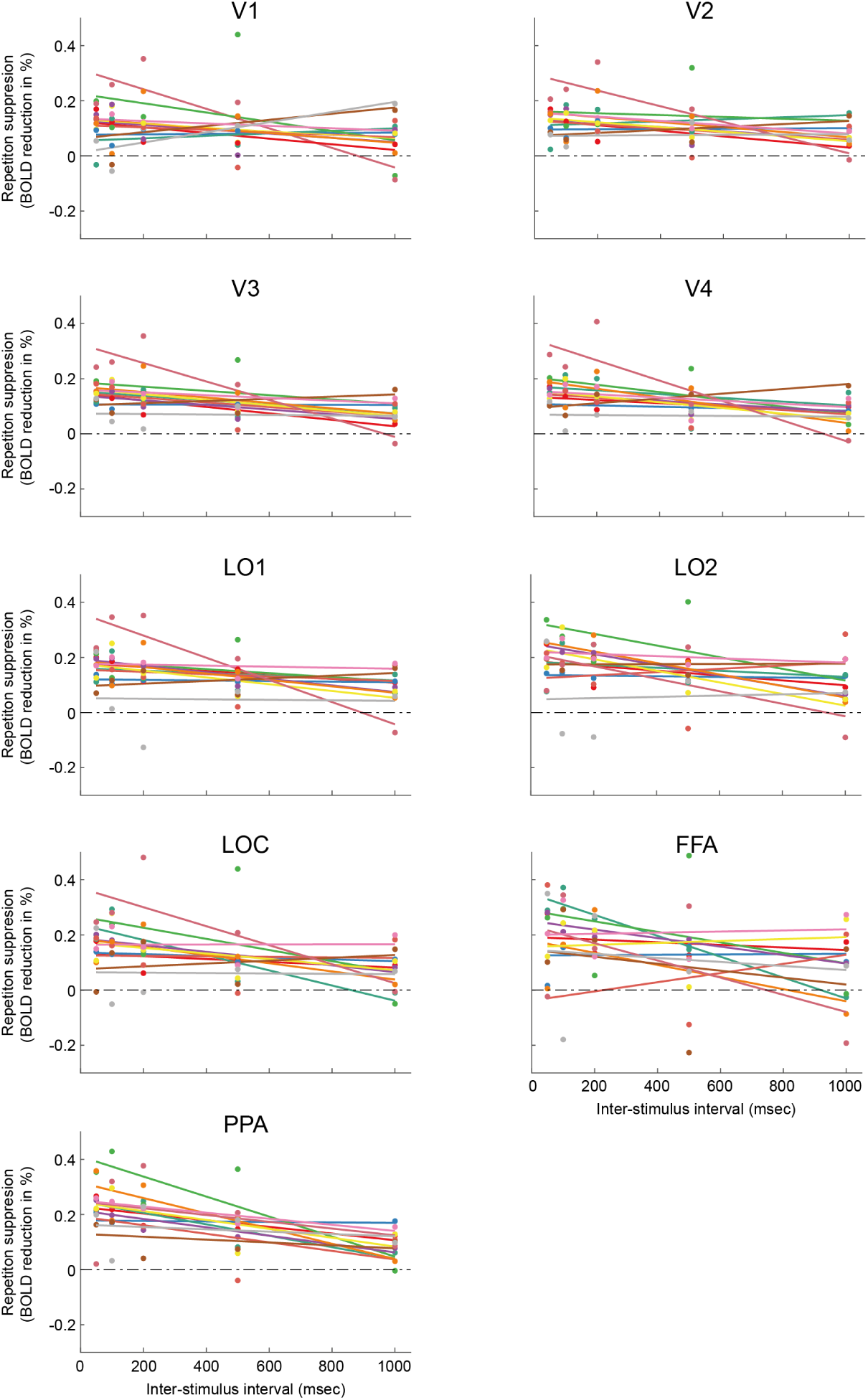
Single-subject repetition suppression (RS) score of each ROI as a function of inter-timulus interval. Colored lines indicate the best fitting linear model for each subject. For group averaged RS scores see Fig 3A.

**Supplemental Fig. S5.**
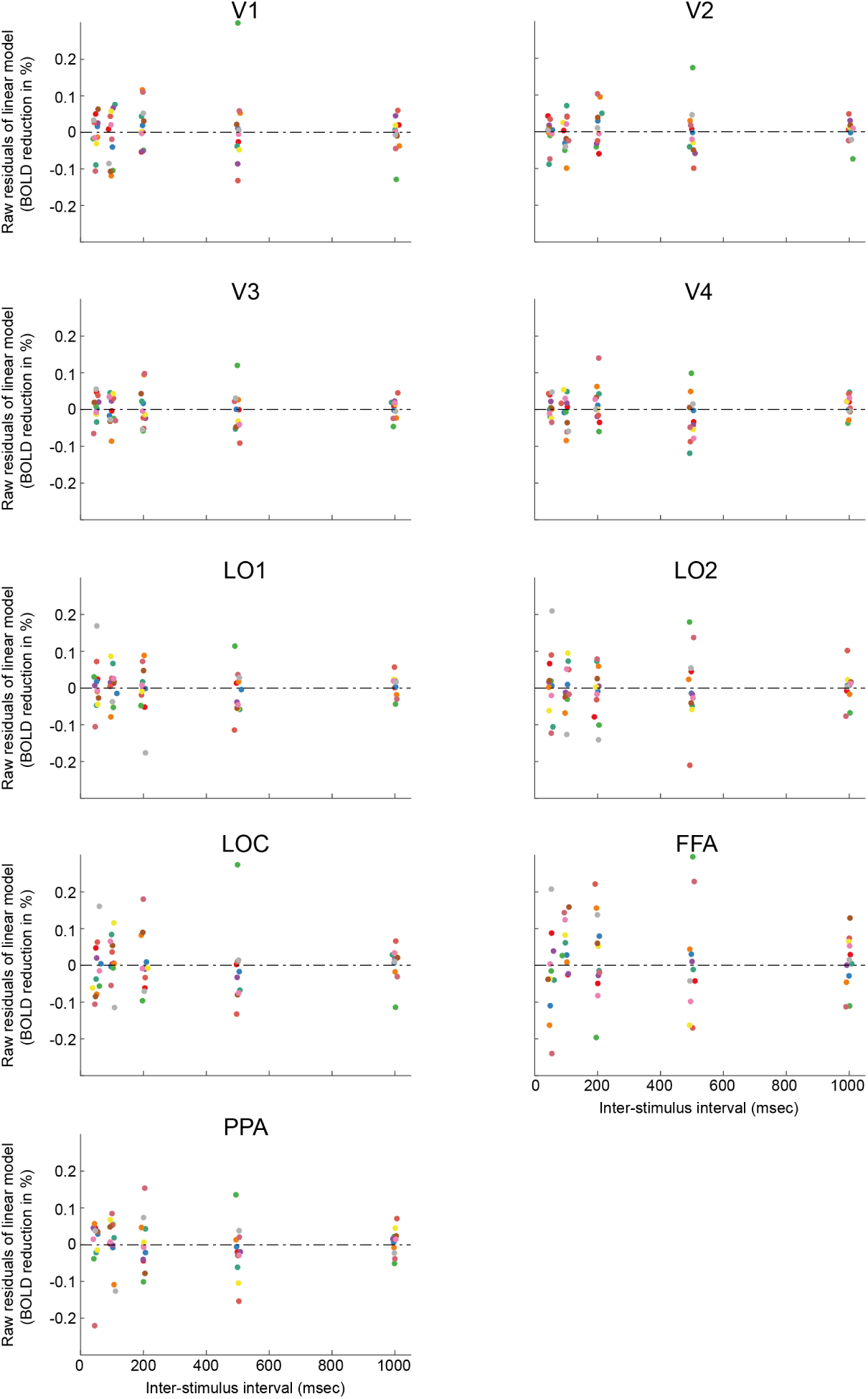
Residuals of linear models Data points depict single-subject deviations from predicted repetition suppression effect There appear to be no systematic deviations from normality. For single-subject RS effect and models see Supplemental Fig S3 and S4

